# Constraints of vigilance-dependent noradrenergic signaling to mouse cerebellar Bergmann glia

**DOI:** 10.1101/2022.06.27.497789

**Authors:** Angelica Salinas-Birt, Xiangyu Zhu, Eunice Y. Lim, Aryana J. Cruz Santory, Liang Ye, Martin Paukert

## Abstract

Behavioral state plays an important role in determining astroglia Ca^2+^ signaling. In particular, locomotion-mediated elevated vigilance has been found to trigger norepinephrine-dependent whole cell Ca^2+^ elevations in astroglia throughout the brain. For cerebellar Bergmann glia it has recently been found that locomotion-induced transient Ca^2+^ elevations depend on their α_1A_-adrenergic receptors. With increasing availability and implementation of locomotion as behavioral parameter it becomes important to understand the constraints of noradrenergic signaling to astroglia. Here we evaluated the effect of speed, duration and interval of locomotion on Ca^2+^ signals in Bergmann glia as well as cerebellar noradrenergic axon terminals. We found almost no dependence on locomotion speed, but following the initial Ca^2+^ transient prolonged locomotion events revealed a steady-state Ca^2+^ elevation. Comparison of time course and recovery of transient Bergmann glia and noradrenergic terminal Ca^2+^ dynamics suggested that noradrenergic terminal Ca^2+^ activity determines Bergmann glia Ca^2+^ activation and does not require noradrenergic receptor desensitization to account for attenuation during prolonged locomotion. Further, analyzing the correlation among Ca^2+^ dynamics within regions within the field of observation we found that coordinated activity among noradrenergic terminals accounts for fluctuations of steady-state Bergmann glia Ca^2+^ activity. Together, our findings will help to better understand astroglia Ca^2+^ dynamics during less controlled awake behavior and may guide the identification of behavioral contexts preferably dependent on astroglia Ca^2+^ signaling.

## 1 INTRODUCTION

There has been an increased focus in the field of neuroscience on the role of astroglia in neuronal circuit function. Chemogenetic manipulations of G protein-coupled receptor activity in astrocytes, which directly affect Ca^2+^ dynamics (Durkee et al., 2019), have revealed insight into potential roles of astrocytes in cognitive function, including learning and memory (Adamsky et al., 2018; Kol et al., 2020; Nagai et al., 2021) and homeostasis of general behavioral activity (Nagai et al., 2019).

Paradigms currently in use to cause and investigate endogenous brain-wide changes in Ca^2+^ activity in astroglia such as spherical or linear treadmills are frequently being used to monitor and stimulate mice during quiet wakefulness or during either spontaneous or motor-enforced locomotion. Monitoring changes in intracellular Ca^2+^ dynamics in awake behaving mice walking on a treadmill provides a way to understand the behavioral context of astroglia Ca^2+^ activity in a variety of brain regions, including the primary visual cortex, somatosensory cortex, and cerebellum (Dombeck et al., 2007; Nimmerjahn et al., 2009; Paukert et al., 2014). Of particular interest are global, whole-cell population Ca^2+^ elevations seen when animals undergo changes in locomotion or are exposed to other stimuli that trigger heightened vigilance (Nimmerjahn et al., 2009; Ding et al., 2013; Srinivasan et al., 2015). These Ca^2+^ elevations occur simultaneously in multiple brain regions (Paukert et al., 2014; Gray et al., 2021). In contrast to these global events, during sleep, Ca^2+^ dynamics are reduced to the microdomain level (Bojarskaite et al., 2020; Ingiosi et al., 2020; Vaidyanathan et al., 2021). In Bergmann glia of the cerebellum and in cortical astrocytes, the population Ca^2+^ elevations are abolished under isoflurane anesthesia, an example of state-dependent activity that demonstrates the importance of examining astroglia Ca^2+^ dynamics in awake animals (Nimmerjahn et al., 2009; Thrane et al., 2012). In accord with reduced locus coeruleus activity during sleep and general anesthesia (Vazey and Aston-Jones, 2014; Hayat et al., 2020) and enhanced locus coeruleus activity during locomotion (Polack et al., 2013; Reimer et al., 2016), these global vigilance-dependent Ca^2+^ responses are norepinephrine-dependent via α_1_-adrenergic receptors (Ding et al., 2013; Paukert et al., 2014; Srinivasan et al., 2015), and in cerebellar Bergmann glia, almost exclusively through their α_1A_-adrenergic receptors (Ye et al., 2020).

Exposing mice to physiological sensory or aversive stimuli such as whisker stimulation, aversive airpuff, or acute isoflurane exposure can cause arousal and increases in astroglia Ca^2+^ activity as seen with locomotor activity (Ding et al., 2013; Srinivasan et al., 2015; Oe et al., 2020; Zuend et al., 2020). Astrocytes in the primary visual cortex are also known to demonstrate potentiated global Ca^2+^ activation when exposed to visual stimulation (Paukert et al., 2014; Slezak et al., 2019) due to enhanced norepinephrine release (Gray et al., 2021). Remarkably, despite the dependence of heightened vigilance-induced astroglia Ca^2+^ elevations on G_q_ protein-coupled α_1_-adrenergic receptors, activation of G_q_ protein-coupled Designer Receptors Exclusively Activated by Designer Drugs (DREADD) overexpressed in cortical astrocytes, suppress astrocyte Ca^2+^ dynamics rendering these cells unresponsive to endogenous receptor activation (Vaidyanathan et al., 2021). Thus, it is important to understand the behavioral constraints that control heightened vigilance-dependent astroglia Ca^2+^ activity.

In this study we employed a linear motorized treadmill to allow for precise controls of both speed and duration during locomotor activity. Astroglia Ca^2+^ responses during enforced locomotion are indistinguishable from responses during spontaneous locomotion events, while the reproducibility of motorized locomotion stimuli facilitates quantification (Paukert et al., 2014). We found that continuous locomotion maintained a sustained component of Bergmann glia Ca^2+^ fluctuations that were not limited by Bergmann glia responsiveness but instead by Ca^2+^ activity in noradrenergic terminals and therefore represent a sustained state of heightened vigilance. Ca^2+^ fluctuations in noradrenergic neurons during extended states of heightened vigilance were coordinated across terminals. Our findings suggest that downstream consequences of vigilance-dependent Bergmann glia Ca^2+^ activation should not be limited to the onset of active behavioral states.

## 2 MATERIALS AND METHODS

### 2.1 Experimental design

All animal procedures were conducted in accordance to guidelines and protocols of the University of Texas Health Science Center at San Antonio (UTHSCSA) Institutional Animal Care and Use Committee. At the time of data collection, all mice were 2-6 months old. For all awake behavior experiments, we employed a previously reported head-restrained mouse on motorized treadmill paradigm. To minimize risk of experimenter bias, all locomotion protocols and subsequent data analysis routines described below were automated. In global *Adra1a* knockout experiments, the experimenter was uninformed of the mouse genotype. On the basis of availability, male and female mice were assigned randomly for all experiments. Statistical analysis was based on number of mice.

### 2.2 Animals

Mice were kept in the Laboratory Animal Resources facility with the ambient temperature maintained at 72–78°F and 30-70% humidity. Mice had *ad libitum* access to water and chow and were maintained on a reverse 12 h-light/12 h-dark schedule (lights off at 9 a.m., on at 9 p.m.). All experimental procedures were performed during the dark cycle. The transgenic mouse breeding strategy resulted in either genetically-encoded Ca^2+^ indicator (GECI) GCaMP6f (Ai95) or Lck-GCaMP6f expressed in a Cre-dependent manner. Each mouse was heterozygous for one GECI allele and heterozygous for one Cre recombinase allele. For global *Adra1a* knockout experiments, experimental mice were heterozygous for GECI, heterozygous for Cre recombinase using the *Slc1a3*-*CreER*^*T*^ mouse line, and homozygous for either the mutant or wt *Adra1a* allele. *Adra1a* mutant homozygous mice appeared healthy and were born in a frequency expected from Mendelian principles. Genotypes were determined from DNA extracted from a toe or tail sample via PCR using primers listed in Supplementary Table 1.

### 2.3 Tamoxifen administration and recombination efficiency

Tamoxifen (Sigma-Aldrich, #T5648-5G) was dissolved in sunflower seed oil (Sigma-Aldrich, #S5007) at a concentration of 10 mg/ml by vortexing and sonication for approximately 30 min and was stored at 4°C for no more than five days. For experiments using the *Slc1a3*-*CreER*^*T*^ mouse line to overexpress GCaMP6f, tamoxifen was injected intraperitoneally at 10 μL/g mouse body weight, starting at age 3–4 weeks, and three times within 5 days. This resulted in GCaMP6f overexpression in all Bergmann glia. For experiments using the *Dbh*-*Cre* mouse line to overexpress Lck-GCaMP6f in noradrenergic neurons, no tamoxifen administration was necessary. Surgeries were done 1–2 weeks after the last tamoxifen injection.

### 2.4 Animal surgery

The surgery procedure had two steps. In the first-step surgery, under intraperitoneal anesthesia (100 mg/kg ketamine and 10 mg/kg xylazine in 0.9% saline solution), the mouse was kept on a heating pad to maintain body temperature at ∼36°C. Following removal of hair above the skull, the skin was disinfected using povidone-iodine. Skin and muscles above the cranium were excised, and 3% hydrogen peroxide was applied to disinfect and prevent bleeding. After scraping away the periosteum, approximately 3mm of muscle surrounding the exposed skull was covered with cyanoacrylate cement. A custom-designed stainless-steel head-plate with a 4mm x 6mm oval opening was centered above lobulus simplex/crus I of the cerebellar hemisphere and then mounted on the skull with dental cement (C&B Metabond, Parkell Inc., Brentwood). While mice recovered from anesthesia, wound edges were coated with Neosporin^®^ ointment containing pramoxine, and mice were placed in a heated cage for recovery. In the second-step surgery, under isoflurane anesthesia (1.5–2% vol./vol. isoflurane in O_2_ with flow rate adjusted based on paw pinch reflex) and on a heating pad, a craniotomy was performed where an area of skull and underlying dura mater of 2 mm × 2 mm were removed and replaced with a glass window comprised of three layers of trimmed No. 1 Corning cover glass (Corning 2975-223) fused with UV-curable Norland optical adhesive 81 (Norland NOA81). The edges of the glass window were sealed with dental cement (Ortho-Jet-Acrylic-Powder, Lang). After one week of post-surgery recovery, mice were habituated to the linear treadmill and the recording conditions in at least three sessions. Imaging was performed at least two weeks after surgeries.

### 2.5 *In vivo* 2P imaging

Two setups were used for Ca^2+^ imaging experiments. For all chronic astroglia *in vivo* experiments, a galvanometer-based 2P laser-scanning Movable Objective Microscope (MOM) (Sutter Instruments) with a x16, 0.80 NA water-immersion objective (Nikon) was used. A pulsed Ti:Sapphire laser beam at 920nm, 80 MHz repetition rate, 140 fs pulse width (Coherent Inc., Chameleon Ultra II) was focused at ∼60 μm below the pial surface reaching halfway through the cerebellar molecular layer. To avoid excessive brain injury, the laser power was attenuated to 12–35 mW at the front aperture of the objective. Emitted light was detected using a gallium arsenide phosphide (GaAsP) photomultiplier tube (H10770PA-40; Hamamatsu Photonics). The awake mouse was positioned on a custom-made linear treadmill, and the head-plate was mounted under the microscope objective. The speed of the treadmill belt was monitored with an optical encoder. The belt was freely movable so that the mouse could walk voluntarily. To enforce locomotion, a servo motor was engaged to move the treadmill at 80–110 mm/s for kinetics and recovery experiments or at the programed speed for the speed experiment. Electromyography (EMG) signals were recorded as the surface potential difference between two silver wires inserted subcutaneously at the right shoulder and left hip of the mouse. The 2P microscope was controlled by an Xi Computer Corporation personal computer (Intel(R) Core(TM) i7-4930 CPU @ 3.40 GHz, 8 GB of RAM) running ScanImage (v3.8.1; Vidrio Technologies, LLC) software within MATLAB R2011b (MathWorks). Image acquisition was triggered at a rate of 1.5 frames/s, and locomotion speed data, EMG data, and Y-mirror position data were simultaneously acquired at 20 kHz sampling rate using National Instruments boards controlled by custom-written scripts in Labview2013 (version 13.0.1f2, National Instruments). Acquired frames were 400 μm × 400 μm at a resolution of 512 pixels per line and 512 lines per frame. Non-imaging data were post hoc downsampled to the image acquisition frame rate and the Y-mirror signal was used to assign appropriate data bins to individual image frames. The entire experimental setup was enclosed in a blackout box.

For experiments on noradrenergic terminal Ca^2+^ dynamics, a resonant scanning version of the MOM (Sutter Instruments) was used with a x16, 0.80 NA water-immersion objective (Nikon). A pulsed Ti:Sapphire laser beam at 920 nm, 80 MHz repetition rate, <120 fs pulse width (Insight DS+, Spectra-Physics MKS Instruments Light & Motion) was used for 2P excitation. The acquisition rate was 15 frames/s with the laser power adjusted to 15–35 mW at the front aperture of the objective. The microscope was controlled by an Xi Computer Corporation personal computer (Intel(R) Core(TM) i7-5930K CPU @ 3.50 GHz, 16 GB of RAM) running ScanImage (v5.0; Vidrio Technologies, LLC) software within MATLAB R2016a (Mathworks). Acquired frames were 400 μm x 400 μm or 100 μm x 100 μm at a resolution of 512 pixels per line and 512 lines per frame. All other design principles including the treadmill were matched between microscopes.

### 2.6 Kinetics and recovery experiments

To measure Bergmann glia and noradrenergic terminal Ca^2+^ dynamics during different durations of locomotion, the Labview data acquisition program called different Arduino board subroutines to control treadmill movement such that enforced locomotion started at a fixed time point and continued for the following periods (s): 1, 2, 5, 10, 30, 60, or 120. Each experiment consisted of three consecutive series of these seven durations of locomotion, and each series proceeded in alternating ascending or descending order. Each mouse was subjected to two experiments where the second experiment began in the opposite order. This strategy minimized any bias regarding potential receptor desensitization or Ca^2+^ store status due to experimental sequence. It also provided six repetitions per locomotion duration, better ensuring at least one repetition would be available for analysis after removal of repetitions where voluntary movement produced Ca^2+^ transients during the baseline period, which attenuates the amplitude of the Ca^2+^ response at the onset of enforced locomotion.

To measure the dynamics of Bergmann glia and noradrenergic terminal Ca^2+^ recovery, mice underwent two 30 s bouts of locomotion with an intermediate pause of varying length (s): 1, 2, 5, 10, 30, 60, or 120. As in the kinetics experiments, each experiment consisted of three consecutive series of these seven durations of intermediate pauses in alternating ascending or descending order. Each mouse was subjected to two experiments where the second experiment began in the opposite order. To analyze the recovery dynamics of Bergmann glia and noradrenergic Ca^2+^ signals, an exponential curve fit was applied to the scatterplot of mean Ca^2+^ changes normalized to the response to the respective first bout of locomotion over specified time points and were fitted using the MATLAB curve fitting application. The custom equation ‘peak response-peak response*exp(-time/time constant)+offset’ was used with the following fit parameters (these parameters were optimized by viewer’s eye in order to best match the datapoints in the majority of plots and were then applied to all fits): method – nonlinear least squares, robust – bisquare, algorithm – Levenberg-Marquardt, MaxFunEvals = 5000, MaxIter = 1000, StartPoint = [0.1 0.1 0.1], DiffMinChange = 0.1, DiffMaxChange = 1, TolFun = 0.5, and TolX = 1.0e-6.

### 2.7 Speed experiment

To determine the effect of speed on Bergmann glia Ca^2+^ dynamics, enforced locomotion occurred for 30 s at the following speeds (mm/s): 10.7, 31.2, 51.4, 70.3, 88.1, 107.9, or 133.6. Each experiment consisted of a single set of these seven speeds in ascending or descending order. Each mouse was subjected to two experiments where the second experiment began in the opposite order.

### 2.8 Data analysis

#### 2.8.1 Locomotion-induced Ca^2+^ dynamics

Imaging data were saved in ScanImage as tiff files, imported to MATLAB R2016a or R2018b, and stored as mat files. Analysis was conducted using custom-written scripts derived from a combination of built-in, open-source, and custom-written functions. Any computer code can be shared upon request. Images were first passed through a Gaussian filter (1.52 SD per pixel distance) to attenuate random noise of the detector. Individual frames within the entire time series were registered to maximize correlation. Since individual structure elements of Bergmann glia or noradrenergic terminal processes cannot be assigned to individual cells, regions of interest (ROIs) were spatially defined by an 8 × 8 checkerboard pattern, which resulted in ROIs of approximately 50 μm x 50 μm. For the few noradrenergic terminal experiments acquired at 100 μm x 100 μm, ROIs were spatially defined by a 2 × 2 checkerboard pattern to preserve the 50 μm x 50 μm ROI size.

Every two consecutive image frames of noradrenergic terminal data were averaged for noise reduction, resulting in a final effective frame rate of 7.5 Hz. For each trial and ROI, the following analysis was performed. The average detector offset was determined by imaging under identical conditions with the laser shutter closed and then subtracted. Median fluorescence value during baseline (from start of a trial until the frame before onset of locomotion) was calculated and then used to calculate the Δ*F*/*F* fluorescence values for each image frame by (*F* − *F*_median_)/*F*_median_, with *F* being the absolute mean fluorescence value within a given ROI in a given image frame. We previously found that voluntary locomotion-induced Bergmann glia Ca^2+^ elevations during the baseline episode can suppress enforced locomotion-induced responses. Therefore, we excluded those repetitions where at least two consecutive Δ*F*/*F* values during baseline exceeded 3x the average standard deviation (SD) of baseline Δ*F*/*F* values of all trials of a data set. Ca^2+^ responses within all individual ROIs of a field of view were analyzed first and population data in all figures represent the median of all respective ROIs of a mouse. The term “mean Ca^2+^ change” represents mean Δ*F*/*F* within 12 s from onset of locomotion. “Time to peak” (TTP) represents the time from onset of locomotion to the maximum Δ*F*/*F* value within 12 s from onset of locomotion. The “correlation coefficient” represents the mean Pearson’s linear correlation coefficient among selected ROI Δ*F*/*F* traces within 20 s from onset of locomotion averaged across three runs of analysis as follows: To compare correlation among ROIs within locomotion trials versus across locomotion trials in an unbiased way, we first pseudo-randomly (each ROI represented at most once) determined a set of 12 ROIs for each “kinetics” experiment described above. Each “kinetics” experiment contained three trials of 120 s locomotion. To analyze “within trial” correlation we determined the mean of all pairwise correlations among the12 ROIs separately for each trial and then averaged the results from all three trials. To analyze “across trial” correlation we divided the set of 12 ROIs into three groups of ROI quadruples. Mean of all pairwise correlations among ROI responses of the first quadruple during the first trial, of the second quadruple during the second trial and of the third quadruple during the third trial was determined. This analysis was repeated for the second quadruple during the first trial, the third quadruple during the second trial and the first quadruple during the third trial. Finally, it was repeated for the third quadruple during the first trial, the first quadruple during the second trial and the second quadruple during the third trial. The average of these three across trials correlation analyses was represented as “across trial” correlation.

#### 2.8.2 Statistical analysis

Statistical analysis was performed using MATLAB R2016a or R2018b (MathWorks). For every group in a data set, the Lilliefors test was applied to test for Gaussian distribution. If all groups followed a Gaussian distribution, parametric statistical tests were applied. If two unrelated normally distributed groups were compared, unpaired two-tailed Student’s *t*-test was applied. For more than two unrelated normally distributed groups, one-way ANOVA was applied and followed by Tukey-Kramer correction for multiple comparisons. If at least one group did not follow Gaussian distribution, non-parametric statistical tests were applied. If two or more unrelated non-normally distributed groups were compared, the Kruskal–Wallis test was used followed by Tukey-Kramer correction for multiple comparisons. The basis of the sample number for individual tests and respective test type applied are mentioned in the associated figure legend. The significance of test results is indicated by *p* values in the graphs or by the abbreviation n.s. for ‘not significant’ when *p* ≥ 0.05.

## 3 RESULTS

### 3.1 Continuous state of heightened vigilance is accompanied by sustained Ca^2+^ fluctuations in cerebellar Bergmann glia

To systematically investigate vigilance-dependent cerebellar Bergmann glia Ca^2+^ dynamics we used *Slc1a3-CreER*^*T* +/-^;Ai95^+/-^ transgenic mice, which express the cytosolic genetically-encoded Ca^2+^ indicator GCaMP6f (Madisen et al., 2015) in astroglia, including cerebellar Bergmann glia (Paukert et al., 2014). GCaMP6f signals were monitored with two-photon microscopy through a chronic cranial window above lobulus simplex and crus I of the cerebellar hemisphere. The duration of heightened vigilance of head-restrained mice was controlled by enforced locomotion on an intermittently motorized and linear treadmill. The advantage of this approach is that mice are able to walk voluntarily or sit relaxed when the motor is off, with no need to balance a spherical treadmill as has been used alternatively to study behavioral state-dependent Bergmann glia Ca^2+^ dynamics (Nimmerjahn et al., 2009). Motorization of the treadmill enables control over timing and intensity of locomotion, representing a behavioral state-clamp paradigm for more precise quantification of vigilance-dependent signaling processes. For quantification of Bergmann glia Ca^2+^ responses in the molecular layer, 40 - 60 μm from the pial surface, an 8 by 8 grid of regions of interest (ROIs) was applied to the 400 μm by 400 μm field of view (Figure S1). Ca^2+^ responses within all individual ROIs of a field of view were analyzed first and population data in all figures represent the median of all respective ROIs of a mouse. It has previously been found that 5 s bouts of enforced locomotion produce transient Bergmann glia Ca^2+^ elevations indistinguishable from voluntary locomotion events in terms of time course and there is no correlation between voluntary locomotion speed and the size of the Ca^2+^ responses (Paukert et al., 2014). In agreement with the latter finding, we now found that the rise time as well as mean amplitude of transient Bergmann glia Ca^2+^ elevations were indistinguishable in response to a wide range of enforced locomotion speed (50 - 130 mm/s; Figure S2). For consistency, when we use the terms locomotion or vigilance, we employed enforced locomotion within 90 - 110 mm/s throughout this study. Varying the length of locomotion revealed that locomotion events of 5 s or longer led to transient Bergmann glia population Ca^2+^ elevations of saturating mean amplitudes (Figure 1a-c) and the time from onset of locomotion to the peak of the transient response was independent of length of locomotion (Figure 1d).

**Figure 1.**
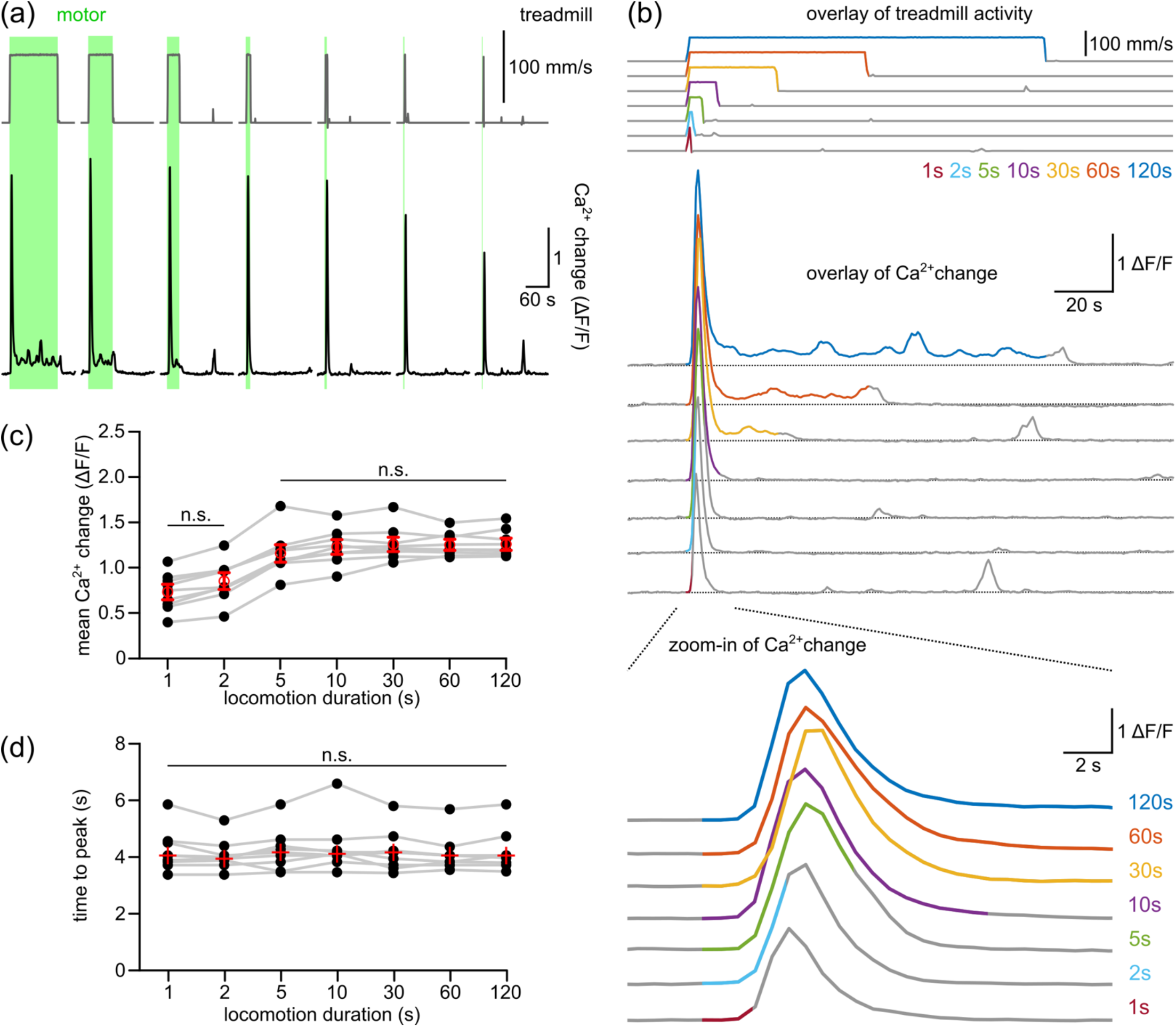
Effects of varying durations of enforced locomotion on vigilance-dependent Bergmann glia Ca^2+^ elevations. (a) Representative example from an awake *Slc1a3-CreER*^*T*^;Ai95 mouse. Upper, gray traces indicate treadmill speed (green bars, enforced locomotion) with different durations of locomotion (from left to right (s): 120, 60, 30, 10, 5, 2 and 1); lower, black traces represent corresponding Bergmann glia Ca^2+^ dynamics. (b) Overlay of stimulus onset-aligned locomotion events (upper) and color-matched Ca^2+^ fluorescence traces in response to different durations of enforced locomotion (lower). (c) Population data in respect to (a), the mean Δ*F/F*_12s_ or the normalized mean Ca^2+^ change ((*F* − *F*_median_ of baseline)/*F*_median_ of baseline) within 12 s from onset of locomotion, during different durations of locomotion. (d) Population data in respect to (a), the mean time to peak (time from onset of locomotion to peak of population response), during different durations of locomotion. Statistical tests used were (c), one-way ANOVA (*F*_6,56_ = 12.345, *p* < .0001; *n* = 9 mice) followed by Tukey-Kramer correction and (d), Friedman test followed by Tukey–Kramer correction (*p* = 0.0814). Red circles with error bars represent mean ± SEM and were used if data follow Gaussian distribution; red crosses without error bars represent median and were used if data do not follow Gaussian distribution. Dots connected by gray lines indicate data from the same mouse. ‘n.s.’ or ‘not significant’ indicates *p* > 0.05. Source data and *p* values are available in source data files.

Following the initial transient Bergmann glia Ca^2+^ elevation, the response attenuated to a sustained phase of apparently stochastic Ca^2+^ fluctuations at a mean amplitude of 9.4% of the initial transient peak amplitude (Figure 1a and b; median, range: 8.2% - 14.6%; *n* = 9 mice).

### 3.2 Sustained vigilance-dependent Ca^2+^ fluctuations in cerebellar Bergmann glia depend on noradrenergic signaling through α_1A_-adrenergic receptors

Transient Bergmann glia Ca^2+^ elevations in response to arousing stimuli such as brief bouts of enforced (5 s) or voluntary locomotion or aversive air puffs depend on α_1A_-adrenergic receptors (Ye et al., 2020). The sustained Ca^2+^ dynamics observed during continuous locomotion could be due to activation of α_1A_-adrenergic receptors or another adrenergic receptor, or it could represent norepinephrine-independent signaling. Gene deletion of α_1A_-adrenergic receptors was sufficient to almost completely abolish the transient and the sustained components of Bergmann glia Ca^2+^ elevations during continuous locomotion (Figure 2). Only a very small sustained Ca^2+^ elevation persisted at 15.0% the size of wildtype littermates (Figure 2b and d), which was 1.5% of the wildtype peak response. These findings suggest that global Bergmann glia Ca^2+^ dynamics represent the vigilance state not only during the startling onset of locomotion but also during prolonged periods of locomotion activity. The considerable attenuation of the intensity of Bergmann glia Ca^2+^ dynamics at the transition from transient to sustained responses could indicate that sustained Ca^2+^ dynamics are limited by desensitization of α_1A_-adrenergic receptors or by Ca^2+^ availability in Bergmann glia internal stores. Each mechanism would limit Bergmann glia Ca^2+^ responsiveness. Alternatively, the excitation of noradrenergic nerve terminals and subsequent release of norepinephrine may determine how much Bergmann glia Ca^2+^ dynamics can be sustained during continuous locomotion.

**Figure 2.**
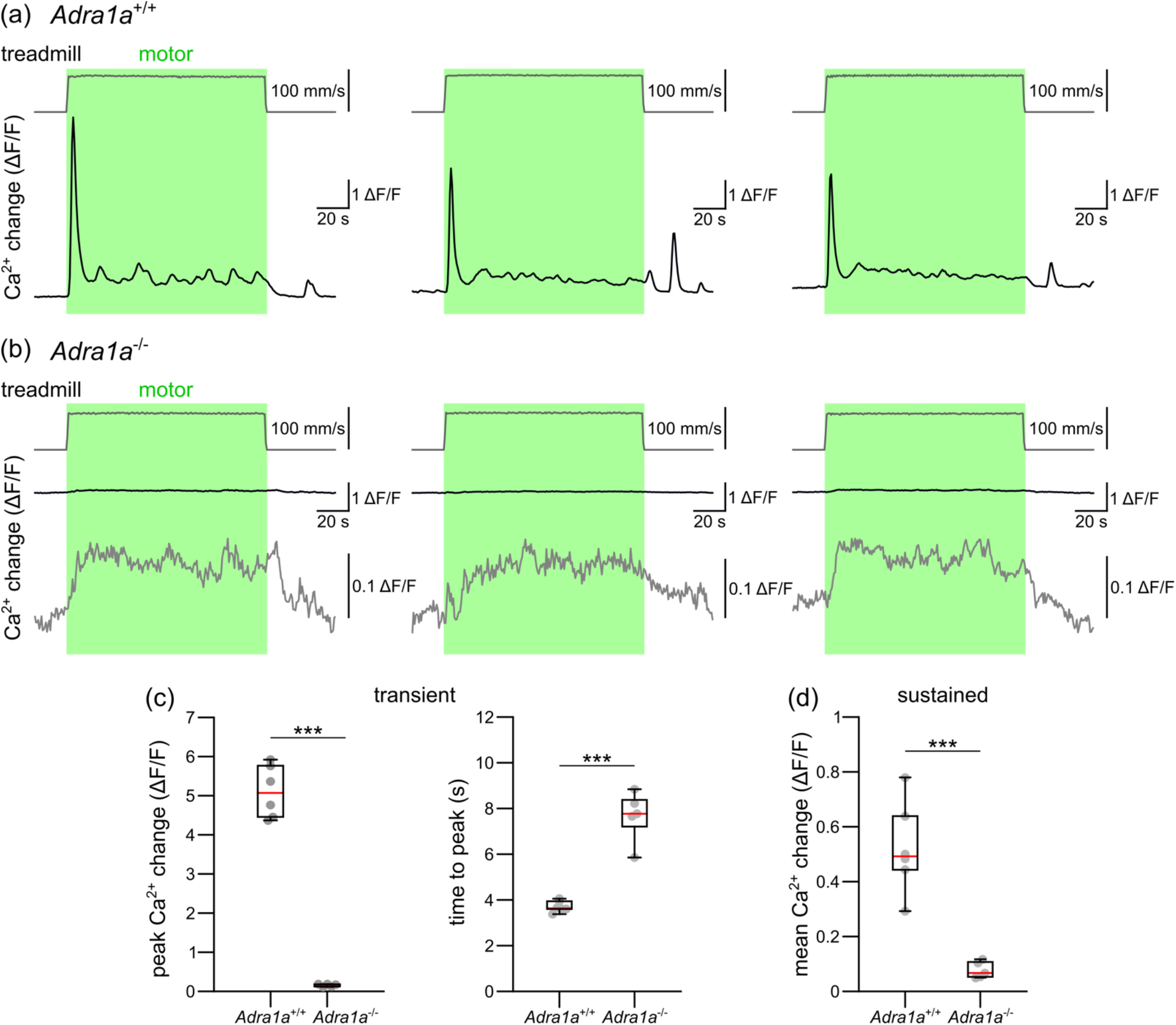
α_1A_-adrenergic receptor is required for sustained vigilance-dependent BG Ca^2+^ activation. (a) Three representative examples from an awake *Slc1a3-CreER*^*T*^;Ai95;*Adra1a*^+/+^ mouse. Upper, gray traces indicate treadmill speed (green bars, 120 s enforced locomotion); lower, black traces represent corresponding Bergmann glia Ca^2+^ dynamics. (b) Three representative example from an awake *Slc1a3-CreER*^*T*^;Ai95;*Adra1a*^-/-^ mouse. Upper, gray traces indicate treadmill speed (green bars, 120 s enforced locomotion); middle, black traces represent corresponding Bergmann glia Ca^2+^ dynamics; lower, same as middle, but with different scale. (c) Left, population data in respect to (a) and (b), the mean Ca^2+^ change within 12 s from onset of locomotion, during different durations of locomotion in *Adra1a*^*+/+*^ and *Adra1a*^*-/-*^ mice. Right, population data in respect to (a) and (b), the mean time to peak during different durations of locomotion in *Adra1a*^*+/+*^ and *Adra1a*^*-/-*^ mice. (d) Population data comparing mean Ca^2+^ change during the sustained portion of locomotion (12^th^ - 120^th^ s from onset of locomotion) in *Adra1a*^*+/+*^ and *Adra1a*^*-/-*^ mice. *Adra1a*^*+/+*^, *n* = 6 mice and *Adra1a*^*-/-*^, *n* = 5 mice. Statistical analysis used were (c, left) unpaired, two-tailed Student’s *t*-test, *t*(9) = 16.372, *p* < 0.001; (c, right) unpaired, two-tailed Student’s *t*-test, *t*(9) = -8.542, *p* < 0.001; and (d) unpaired, two-tailed Student’s *t*-test, *t*(9) = 5.795, *p* < 0.001. Asterisks indicate significant difference (*, *p* < 0.05; **, *p* < 0.01; ***, *p* < 0.001) and ‘n.s.’ or ‘not significant’ indicates *p* > 0.05. Gray dots indicate individual data point from each mouse, overlayed with box and whisker plot with defined elements, median (center line), upper and lower quartiles (bounds of box), and highest and lowest values (whiskers). Source data and *p* values are provided in source data file.

### 3.3 Vigilance-dependent Bergmann glia transient Ca^2+^ elevations recover within tens of seconds of rest resembling the recovery of noradrenergic terminal Ca^2+^ elevations

To assess the recovery of Bergmann glia capability to respond to the onset of locomotion with a transient Ca^2+^ elevation, we used 30 s bouts of enforced locomotion to let Bergmann glia reliably reach the sustained response phase (Figure 3a and b). A second 30s bout of locomotion was then applied following periods of rest for 1 to 120 s. This protocol was applied with increasing periods of rest and with decreasing periods of rest for each mouse (see Methods section for details). Fitting of data from individual mice revealed that Bergmann glia transient Ca^2+^ elevations in response to the onset of locomotion recovered with a time constant of 33.5 ± 8.7 s (Figure 3c; *n* = 7 mice). The considerable range of recovery time constant values across individual experiments within mice (Figure S3a) suggested that Bergmann glia Ca^2+^ elevations might rather be limited by the amount of arousal that a new locomotion event could trigger following a preceding 30 s bout of locomotion than by Bergmann glia responsiveness. To test this possibility, we imaged Ca^2+^ dynamics in cerebellar cortex noradrenergic terminals. Membrane-tethered GCaMP6f was expressed in noradrenergic neurons by crossing *Dbh-Cre* (Gerfen et al., 2013) and *Lck-GCaMP6f* ^flox^ (Srinivasan et al., 2016) mice. At GCaMP6f expression levels achieved using transgenic mice, membrane-tethering of GCaMP6f is needed to detect noradrenergic terminal Ca^2+^ dynamics (Ye et al., 2020). Since membrane-tethering of GCaMP6f does not allow to visualize the morphology of individual axons *in vivo* (Ye et al., 2020; Gray et al., 2021) we employed the same unbiased 8 by 8 grid of ROIs strategy for analysis of Ca^2+^ responses as described for Bergmann glia (Figure S4). Noradrenergic terminals responded to 30 s bouts of continuous locomotion with transient Ca^2+^ elevations (Figure 4a and b). Using the same protocol described for Bergmann glia, analysis of the recovery of noradrenergic terminals’ transient Ca^2+^ responses to the onset of locomotion revealed that noradrenergic terminal Ca^2+^ responses recovered similarly fast as Bergmann glia responses (Figure 4c and d, Figure S3b; 37.8 ± 4.4 s; *t*(11) = -0.419, *p* = 0.684; *n* = 6 mice; unpaired, two-tailed Student’s *t*-test). These findings are consistent with a model where transient Bergmann glia Ca^2+^ elevations are limited by arousal or the vigilance state of the mouse.

**Figure 3.**
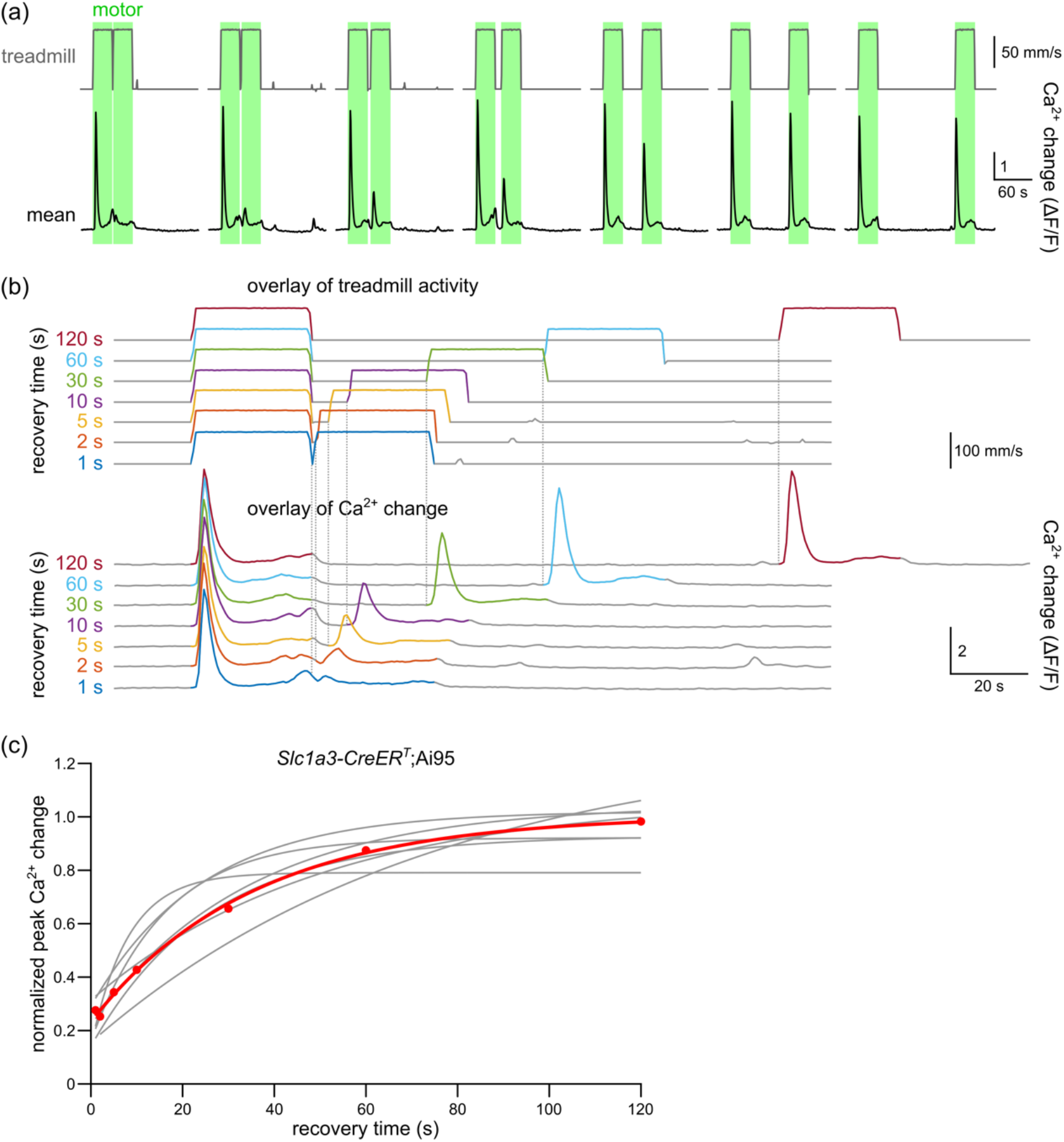
Effect of varying durations of recovery between bouts of enforced locomotion on vigilance-dependent Bergmann glia Ca^2+^ elevations. (a) Representative example from an awake *Slc1a3-CreER*^*T*^;Ai95 mouse. Upper, gray traces indicate the speed of locomotion (green bars, enforced locomotion) with varying lengths of an intermediate pause between two 30 s bouts of locomotion (from left to right in seconds: 1, 2, 5, 10, 30, 60 and 120 s); lower, black traces represent corresponding Bergmann glia Ca^2+^ dynamics. (b) Overlay of stimulus onset-aligned locomotion events (upper) and color-matched Ca^2+^ fluorescence traces (lower) in response to enforced locomotion with different intervening pause lengths. (c) Exponential fitted curves of peak Bergmann glia Ca^2+^ change during second bout of locomotion normalized to peak Ca^2+^ change during the first bout. Gray traces represent the fitted curves of individual mice; red dots represent the median of all mice’s normalized Ca^2+^ change for each respective time point, and the red line indicates the fitted curve of all red dots or median values.

**Figure 4.**
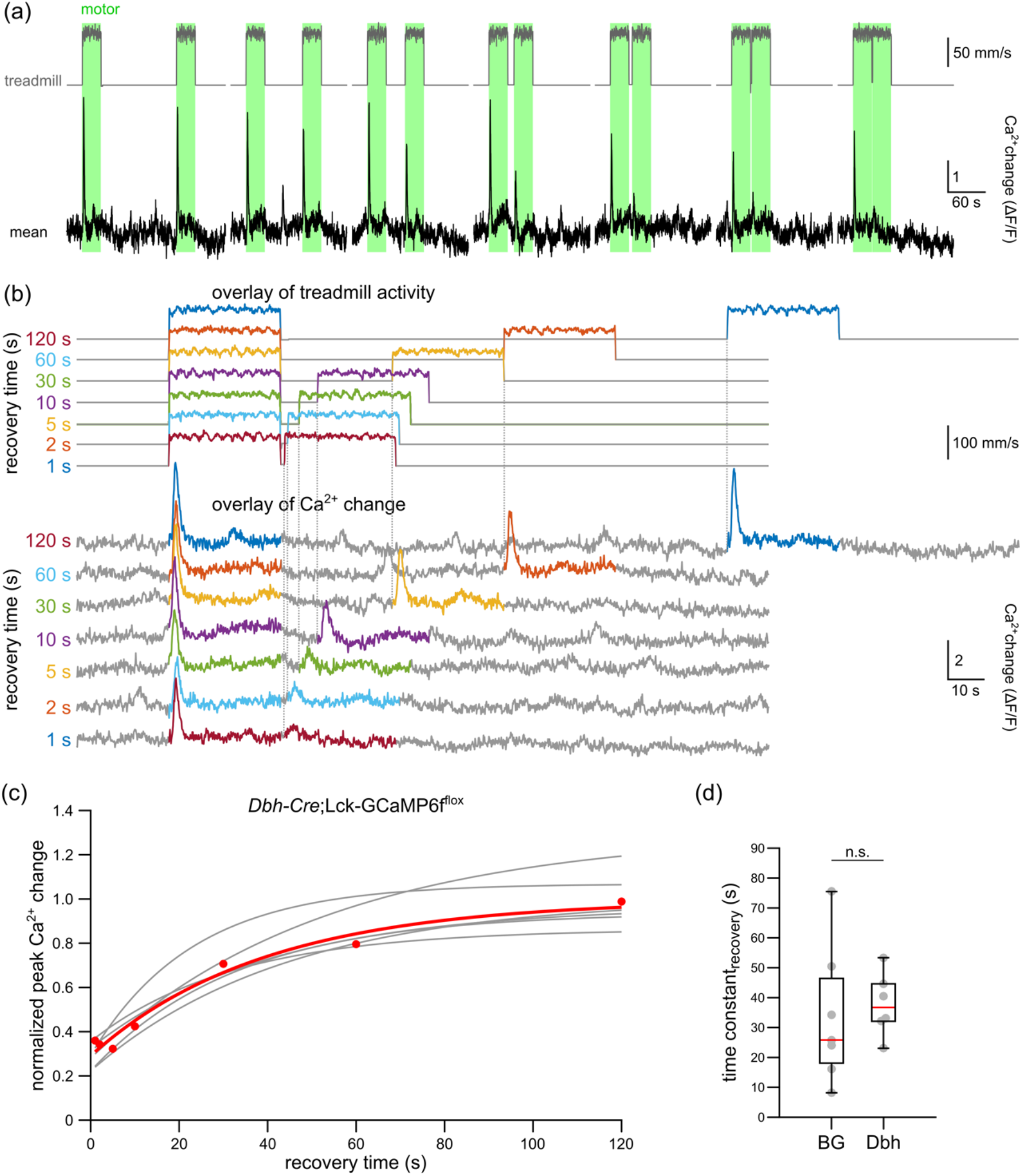
Effect of varying durations of recovery between bouts of enforced locomotion on vigilance-dependent noradrenergic terminals Ca^2+^ elevations. Representative example from an awake *Dbh-Cre*;Lck-GCaMP6f^flox^ mouse. Upper, gray traces indicate the speed of locomotion (green bars, enforced locomotion) with varying lengths of an intermediate pause between two 30 s bouts of locomotion (from left to right (s): 120, 60, 30, 10, 5, 2 and 1); lower, black traces represent corresponding noradrenergic terminals Ca^2+^ dynamics. (b) Overlay of stimulus onset-aligned locomotion events (upper) and color-matched Ca^2+^ fluorescence traces (lower) in response to enforced locomotion with different intervening pause lengths. (c) Exponential fitted curves of peak noradrenergic terminal Ca^2+^ change during second bout of locomotion normalized to peak Ca^2+^ change during the first bout. Gray traces represent the fitted curves of individual mice; red dots represent the median of all mice’s normalized Ca^2+^ change for each respective time point, and the red line indicates the fitted curve of all red dots or median values. (d) Population data of time constant in respect to the fitting in (c). Statistical analysis used unpaired, two-tailed Student’s *t*-test: *t*(11) = -0.419, *p* = 0.684. Gray dots indicate individual data point from each mouse, overlayed with box and whisker plot with defined elements, median (center line), upper and lower quartiles (bounds of box), and highest and lowest values (whiskers). ‘n.s.’ or ‘not significant’ indicates *p* > 0.05. Source data and *p* values are provided as a source data file.

### 3.4 Vigilance-dependent noradrenergic terminal Ca^2+^ elevations precede Bergmann glia responses, are shorter and attenuate at least as much

To further investigate whether locomotion-induced Bergmann glia Ca^2+^ elevations are constrained by noradrenergic terminal Ca^2+^ dynamics we compared the kinetics of these locomotion-induced Ca^2+^ elevations (Figure 5a and b). Noradrenergic terminal transient Ca^2+^ elevations in response to the onset of locomotion reached their peak more than 1 s earlier than Bergmann glia responses (Figure 5a-c). Certainly, the different mechanisms underlying Ca^2+^ elevations in noradrenergic terminals and Bergmann glia need to be considered. N-type Ca^2+^ channels have been found to play a dominant role in release of norepinephrine from sympathetic neurons (Hirning et al., 1988) and therefore may play a similar role in central noradrenergic neurons, while Bergmann glia Ca^2+^ elevations are mediated by G_q_ protein-coupled α_1A_-adrenergic receptors triggered Ca^2+^ release from the endoplasmic reticulum (Ye et al., 2020; Figure 2). Noradrenergic terminal Ca^2+^ elevations also decayed faster, resulting in a shorter Ca^2+^ transient (Figure 5a, b and d) and showed a tendency towards sustaining an even smaller proportion of the respective peak response during continuous locomotion (Figure 5a, b and e). Together, these observations further support a model where transient as well as sustained Bergmann glia Ca^2+^ elevations are limited by arousal or the vigilance state of the mouse.

**Figure 5.**
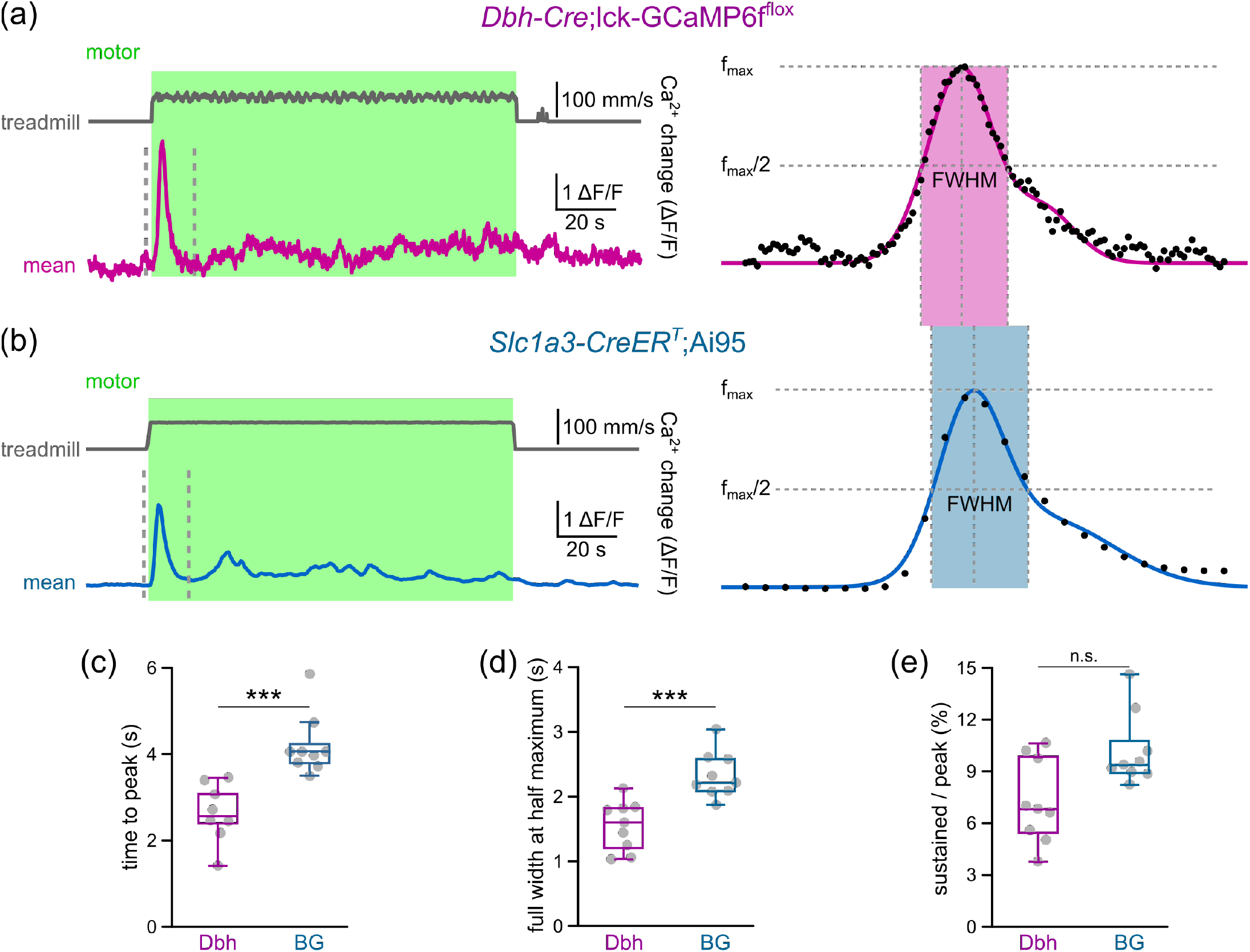
Distinct kinetics of vigilance-dependent Ca^2+^ elevations in noradrenergic terminals and Bergmann glia. (a) Representative example from an awake *Dbh-Cre*;*Lck-GCaMP6f* ^flox^ mouse. Left, gray traces indicate the speed of locomotion (green bars, 120 s enforced locomotion), black traces represent corresponding noradrenergic terminals Ca^2+^ dynamics; right, expanded plots from dotted lines in (a), black dots represent values from individual image frames, magenta bar indicates full width at half maximum (FWHM). (b) Similar as (a), but in *Aldh1l1-CreER*^*T2*^;Ai95 mouse. (c) Population data comparing mean time to peak between noradrenergic terminal Ca^2+^ and Bergmann glia Ca^2+^ dynamics. (d) Population data comparing FWHM. (e) Population data comparing the ratio of the Ca^2+^ amplitude during the sustained phase relative to the amplitude of the preceding peak. *Dbh-Cre*;*Lck-GCaMP6f* ^flox^, *n* = 9 mice and *Aldh1l1-CreER*^*T2*^;Ai95, *n* = 9 mice. Statistical analysis used were (c), unpaired, two-tailed Student’s *t*-test, *t*(16) = 4.493, *p* < 0.001; (d), Kruskal-Wallis Test, *p* < 0.001; and (e), Kruskal-Wallis test, *p* = 0.070. Asterisks indicate significant difference (*, *p* < 0.05; **, *p* < 0.01; ***, *p* < 0.001) and ‘n.s.’ or ‘not significant’ indicates *p* > 0.05. Gray dots indicate individual data point from each mouse, overlayed with box and whisker plot with defined elements, median (center line), upper and lower quartiles (bounds of box), and highest and lowest values (whiskers). Source data are provided as a source data file.

### 3.5 Correlation analysis suggests fluctuating but coordinated norepinephrine release during states of sustained vigilance

Sustained Ca^2+^ dynamics in Bergmann glia as well as noradrenergic terminals during continuous locomotion appeared not only smaller but also more variable than at the onset of locomotion (Figure 6a and b). The availability of norepinephrine-filled vesicles at the release sites closest to the processes of individual Bergmann glia could contribute to variability of their responses, however the small size and variability of noradrenergic terminal sustained Ca^2+^ dynamics suggested that they represented a major constraint of noradrenergic signaling. We then asked whether stochastic Ca^2+^ elevations at individual norepinephrine release sites (could also be individual noradrenergic terminals) accounted for the variability of sustained Ca^2+^ dynamics or whether they represented fluctuations in coordinated activity in at least a cluster of noradrenergic terminals (possibly by global locus coeruleus activation). Observing the sustained Ca^2+^ dynamics in noradrenergic terminals as well as Bergmann glia during two independent trials of continuous locomotion in three respective ROIs covering the entire imaged field of view (Figure 6a and b) revealed examples of apparently coordinated Ca^2+^ fluctuations and also examples of isolated events in both cell types. To systematically investigate whether coordinated activity characterizes noradrenergic signaling to Bergmann glia during extended states of heightened vigilance we pseudorandomly defined sets of 12 ROIs for each imaged field of view. Since each experiment that was described for Figure 1 contained three trials of 120 s continuous locomotion we compared the correlation among sustained Ca^2+^ dynamics of all 12 ROIs during individual trials (“within trials”) with the correlation among the same 12 ROIs when they were separated into three quadruples of ROIs, each quadruple representing data from a different trial (“across trials”; see Methods section for details). We predicted, if stochastic Ca^2+^ elevations at individual norepinephrine release sites or nerve terminals accounted for the variability, the correlation among ROIs should be equally low whether compared within a continuous locomotion trial or compared across different continuous locomotion trials. However, if variability in global locus coeruleus activity or coordinated activity in multiple nerve terminals accounted for the variability in sustained Ca^2+^ dynamics, we would predict increased correlation among ROIs when compared within a continuous locomotion trial as compared to across different continuous locomotion trials. We found a three-fold higher correlation among noradrenergic terminal ROIs within trials than across trials and a two-fold higher correlation among Bergmann glia process ROIs within trials than across trials (Figure 6c). These data indicate that even during continuous locomotion, noradrenergic terminal Ca^2+^ dynamics represent fluctuations of arousal and Bergmann glia Ca^2+^ dynamics reliably report noradrenergic signaling during prolonged states of heightened vigilance.

**Figure 6.**
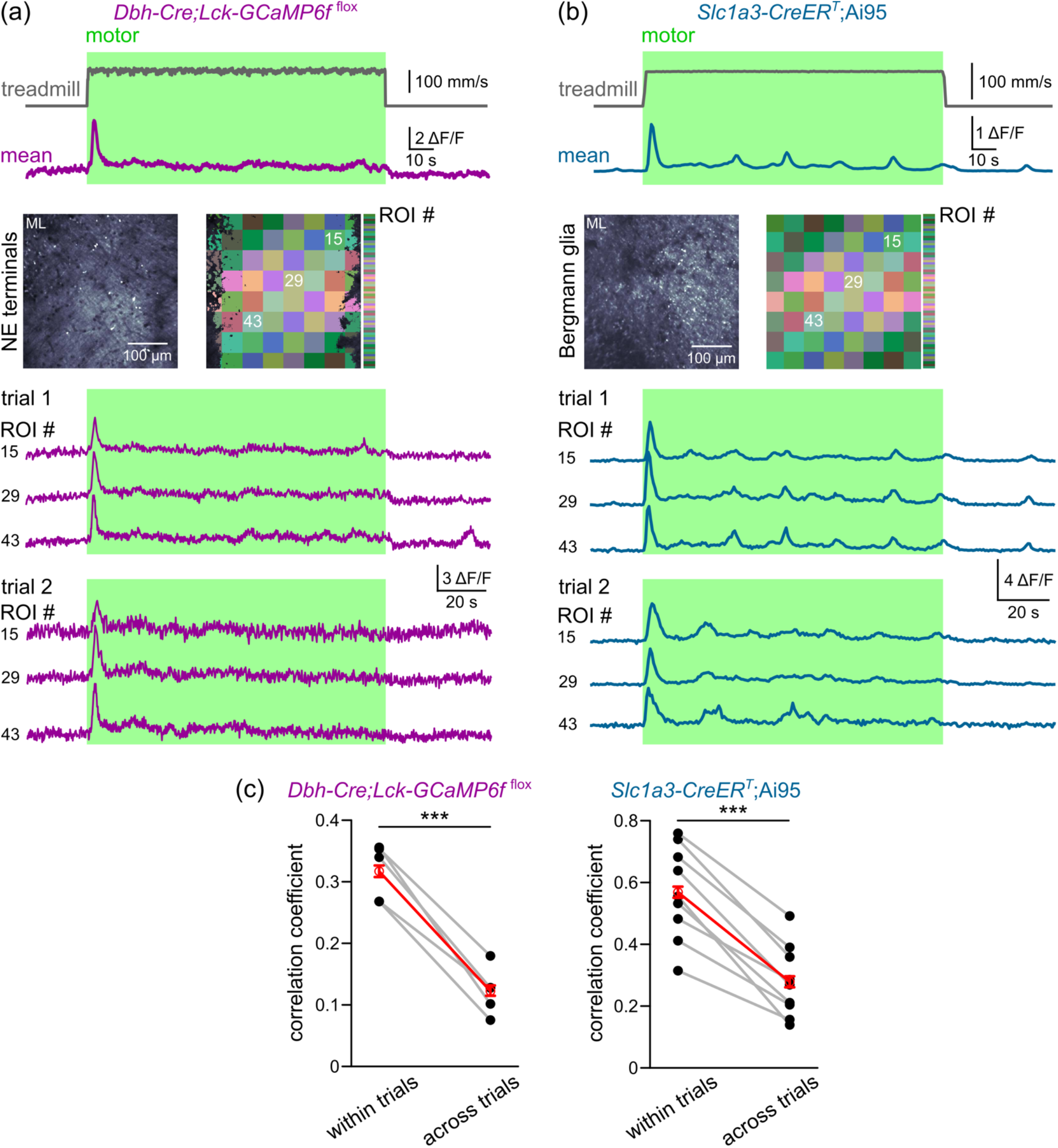
Vigilance-dependent Ca^2+^ elevations in noradrenergic terminals and Bergmann glia correlate with individual enforced locomotion. (a) Representative example from an awake *Dbh-Cre*;*Lck-GCaMP6f* ^flox^ mouse. Upper, gray traces indicate the speed of locomotion (green bars, 120 s enforced locomotion); lower, black traces represent corresponding noradrenergic terminals Ca^2+^ dynamics. Middle, pseudocoloured *in vivo* Ca^2+^ image of noradrenergic terminals and corresponding locations of regions of interest (ROIs). Lower, Ca^2+^ dynamics traces representing numbered ROIs from two representative trials. (b) Similar as (a), but in *Aldh1l1-CreER*^*T2*^;Ai95 mouse. (c) Chained-dot plots compare mean Pearson correlation coefficient of twelve randomly chosen ROIs between intra-events and inter-events in *Dbh-Cre*;*Lck-GCaMP6f* ^flox^ mice (left, *n* = 5 mice) and *Aldh1l1-CreER*^*T2*^;Ai95 mice (right, *n* = 9 mice). Statistical analysis used were (left) paired, two-tailed Student’s *t*-test, *t*(4) = 10.185, p < 0.001; (right), *t*(8) = 9.663, p < 0.001 (*, p < 0.05; **, p < 0.01; ***, p < 0.001). Red symbols represent mean ± SEM, gray lines between dots suggest data from the same mouse. Source data and *p* values are provided in the source data file.

## 4 DISCUSSION

We have taken advantage of control over onset, duration and speed of locomotion with a motorized linear treadmill used conceptually as a behavioral state-clamp. This approach allowed to dissect the constraints of locomotion-induced Bergmann glia global Ca^2+^ elevations and revealed that they reliably represent noradrenergic signaling even during prolonged states of heightened vigilance. This conclusion was supported by gene deletion experiments and by comparing Ca^2+^ dynamics in Bergmann glia processes and noradrenergic terminals in the cerebellar molecular layer during identical locomotion conditions.

Varying the length of locomotion revealed that at the onset of bouts of locomotion lasting 5s or longer, a stereotyped transient and synchronized Ca^2+^ elevation in the entire observed population of Bergmann glia processes was triggered (Figure 1). During a continuous state of heightened vigilance while engaged in locomotion, Bergmann glia Ca^2+^ dynamics were sustained as variable, less synchronized fluctuations with an average amplitude close to 10% of the initial transient peak amplitude. Like the initial transient response, sustained Bergmann glia Ca^2+^ elevations were dependent on α_1A_-adrenergic receptors (Figure 2). The rate and extent of attenuation of the initial transient to sustained Ca^2+^ responses, as well as the recovery of transient responses in Bergmann glia compared to noradrenergic terminals, indicated that noradrenergic terminal excitation rather than Bergmann glia responsiveness constrained noradrenergic signaling (Figures 3-5). Increased correlation between ROIs within trials of continuous locomotion compared to across trials suggested that variable Ca^2+^ fluctuations were due to coordinated excitation of multiple noradrenergic terminals rather than due to independent activity at individual norepinephrine release sites (Figure 6).

The properties of transient Bergmann glia Ca^2+^ elevations at the onset of enforced locomotion were consistent with those previously reported at the onset of voluntary locomotion events. The amplitude of voluntary locomotion-induced Bergmann glia Ca^2+^ elevations does not correlate with locomotion speed (Paukert et al., 2014) as shown here with amplitude and time to peak of responses to a wide range of enforced locomotion speeds (Figure S1). Similarly, Bergmann glia transient Ca^2+^ elevations last longer than short voluntary locomotion events, but decline before longer voluntary locomotion events end (Nimmerjahn et al., 2009). This is consistent with the stereotyped shape of transient Ca^2+^ elevations found at the onset of enforced locomotion for various lengths (Figure 1). There is also a refractory period of 30 s of rest before voluntary locomotion can exert Ca^2+^ elevations of similar size independent of prior locomotion history (Nimmerjahn et al., 2009). This value is in accord with the time constant of recovery of transient Bergmann glia Ca^2+^ elevations at the onset of enforced locomotion (Figure 3). It has been proposed that this refractory period could be caused by depletion of Bergmann glia intracellular stores (Nimmerjahn et al., 2009), and alternatively or in addition, desensitization of α_1A_-adrenergic receptors, which cause locomotion-induced transient intracellular release in Bergmann glia (Ye et al., 2020), could contribute. However, by studying cerebellar molecular layer noradrenergic terminal Ca^2+^ dynamics under equivalent behavioral context we found that they were transient as well, declining faster than Bergmann glia responses (Figure 5), and had recovery kinetics similar to Bergmann glia responses (Figure 4). These findings suggest that excitation of noradrenergic terminals constrains heightened vigilance-dependent Bergmann glia Ca^2+^ activation rather than Bergmann glia responsiveness. The contrast between our finding that Bergmann glia are capable to reliably encode the physiological dynamic range of noradrenergic signaling during various levels of vigilance and the finding that activation of G_q_ protein-coupled DREADDs in astrocytes eventually silences the same signaling pathway (Vaidyanathan et al., 2021) illustrates the importance of studying astroglia Ca^2+^ dynamics through their endogenous receptors activated by behavioral stimuli.

Whereas prolonged (10 s or longer) optogenetic excitation of cortical noradrenergic terminals or arousal associated with fear conditioning is needed to increase the second messenger cAMP in astrocytes, a single air puff to the mouse face was sufficient to cause a transient increase in norepinephrine and a robust astrocyte Ca^2+^ response that did not result in an increase in cAMP (Oe et al., 2020). Consistent with this finding, a 20 s bout of arousal caused by acute exposure to the anesthetic isoflurane causes β-adrenergic receptor mediated glycogenolysis and lactate release from cortical astrocytes in mice (Zuend et al., 2020). Here, combining continuous enforced locomotion as behavioral state-clamp to assess whether locomotion can induce prolonged vigilance with gene deletion revealed that Bergmann glia maintain a sustained component of variable Ca^2+^ fluctuations, which depends on α_1A_-adrenergic receptor activation. Comparing Ca^2+^ dynamics in subsets of ROIs within continuous locomotion trials versus across trials showed a more than twofold higher correlation of Bergmann glia as well as noradrenergic terminal Ca^2+^ dynamics when analyzed within continuous locomotion trials. This finding indicates that variable sustained Ca^2+^ dynamics are often caused by fluctuations of coordinated excitation of groups of noradrenergic terminals, possibly reflecting fluctuations in global locus coeruleus activity, rather than by independent excitation at individual norepinephrine release sites. During quiet wakefulness, noradrenergic terminal GCaMP6f signals did not indicate any activity, which is consistent with prior reports using the same experimental approach in cerebellum or visual cortex (Ye et al., 2020; Gray et al., 2021). In contrast, with virally-delivered expression of cytosolic GCaMP6f in cortical noradrenergic terminals, single-peak as well as multi-peak Ca^2+^ elevations have been found (Oe et al., 2020). It has been suggested that multi-peak events are triggered by bursts of action potentials. Viral delivery is expected to lead to a higher expression level than expression from a single transgene allele in transgenic mice. This may be the reason why in transgenic mice expression of cytosolic GCaMP6f in noradrenergic terminals fails to detect any Ca^2+^ dynamics, and instead membrane-tethering of GCaMP6f is needed (Ye et al., 2020). Hence, Lck-GCaMP6f signals in this study should mainly represent burst firing activity in noradrenergic terminals during heightened vigilance. This interpretation would be consistent with fluctuating coordinated activation of noradrenergic terminals during extended periods of vigilance. The observation that noradrenergic terminal Ca^2+^ dynamics decreased considerably to a sustained level of activity following the initial response at onset of locomotion is consistent with more intense phasic locus coeruleus activation by sensory stimulation during a quiet awake state than during an active awake state (Hayat et al., 2020). Apparently, location and timing of Bergmann glia Ca^2+^ dynamics are important for their functional role. Ca^2+^-permeable α-amino-3-hydroxy-5-methyl-4-isoxazolepropionic acid (AMPA)-type glutamate receptors on Bergmann glia are important for fine motor coordination (Saab et al., 2012) but loss of α_1A_-adrenergic receptors does not impair motor coordination and may rather play a role in the cerebellar cognitive function (Schmahmann, 2019; Ye et al., 2020).

## 5 CONCLUSION

Systematic comparison of locomotion-induced Bergmann glia and noradrenergic terminal Ca^2+^ dynamics in the cerebellar molecular layer revealed that global Bergmann glia population Ca^2+^ dynamics are constrained by noradrenergic terminal activation and therefore reliably represent the current vigilance state. The finding that the dynamic range of noradrenergic signaling to Bergmann glia covers various levels of vigilance during extended periods of behavioral activity should help to better predict the behavioral context where Bergmann glia Ca^2+^ dynamics are needed for normal circuit function and behavioral performance.

## Acknowledgments

We thank Priscilla M. Barba-Escobedo and John Cavaretta for expert support in genotyping and animal husbandry. Eunice Y. Lim was supported by a T32NS082145 fellowship. Aryana J. Cruz Santory was supported by an R25GM095480 fellowship. This work was supported by R01MH113780 and The Robert J. Kleberg, Jr. and Helen C. Kleberg Foundation (Martin Paukert).

