## Supplementary_Materials for "Constraints of vigilance-dependent noradrenergic signaling to mouse cerebellar Bergmann glia"

### SUPPLEMENTARY MATERIAL

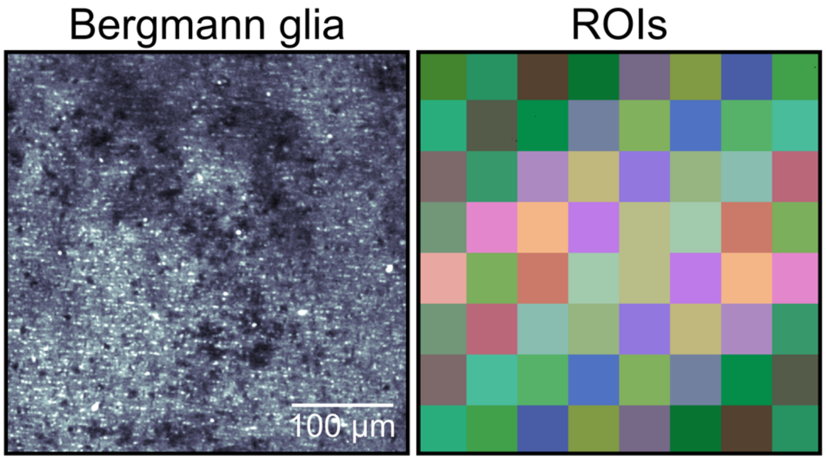


**Supplementary Figure 1**

**Representative field-of-view for a *Slc1a3-CreER$^T$*;Ai95 mouse.**

Image of *in vivo* GCaMP6f in Bergmann glia processes, tangential optical section ~60 µm beneath pial surface (left) and assigned ROIs (right).

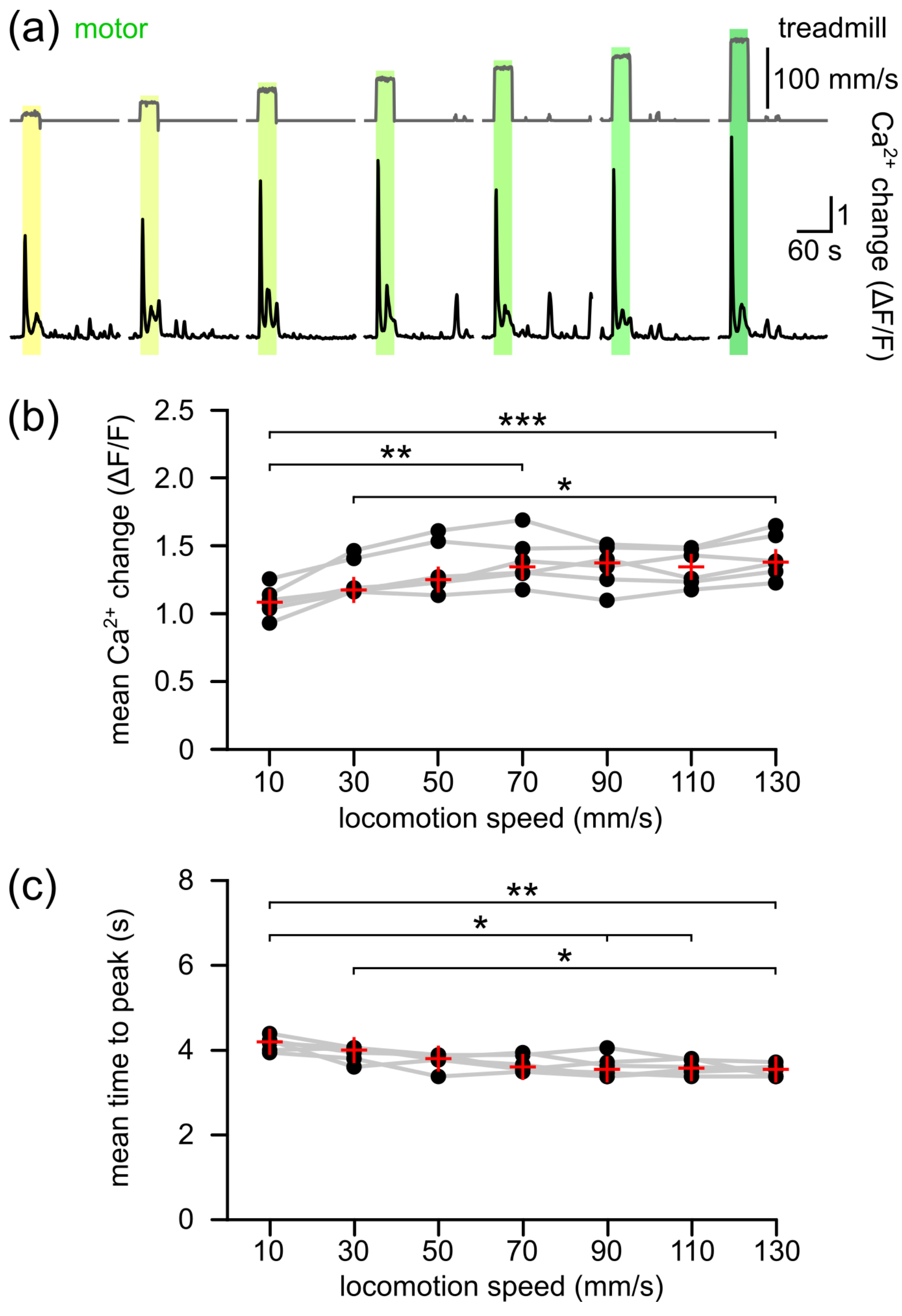


**Supplementary Figure 2**

**Effects of varying speeds of enforced locomotion on vigilance-dependent Bergmann glia $Ca^{2+}$ elevations.**

(a) Representative example from an awake *Slc1a3-CreER*$^{T}$;Ai95 mouse. Upper, gray traces indicate different speeds of enforced locomotion (from left to right (mm/s): 10, 30, 50, 70, 90, 110, and 130); lower, black traces represent corresponding Bergmann glia $Ca^{2+}$ dynamics. (b) Population data in respect to (a), mean $\Delta F/F_{12s}$ during different speeds of enforced locomotion. (c) Population data in respect to (a), mean time to peak during different speeds of enforced locomotion. Statistical tests used were Friedman test followed by Tukey-Kramer correction ($p < 0.001$ for each; $n = 6$ mice). Red crosses without error bars represent median and were used when data do not follow Gaussian distribution. Dots connected by gray lines indicate data from the same mouse. Asterisks indicate significant difference (*, $p < 0.05$; **, $p < 0.01$; ***, $p < 0.001$) and ‘n.s.’ or ‘not significant’ indicates $p > 0.05$. Source data and $p$ values are provided in source data file.

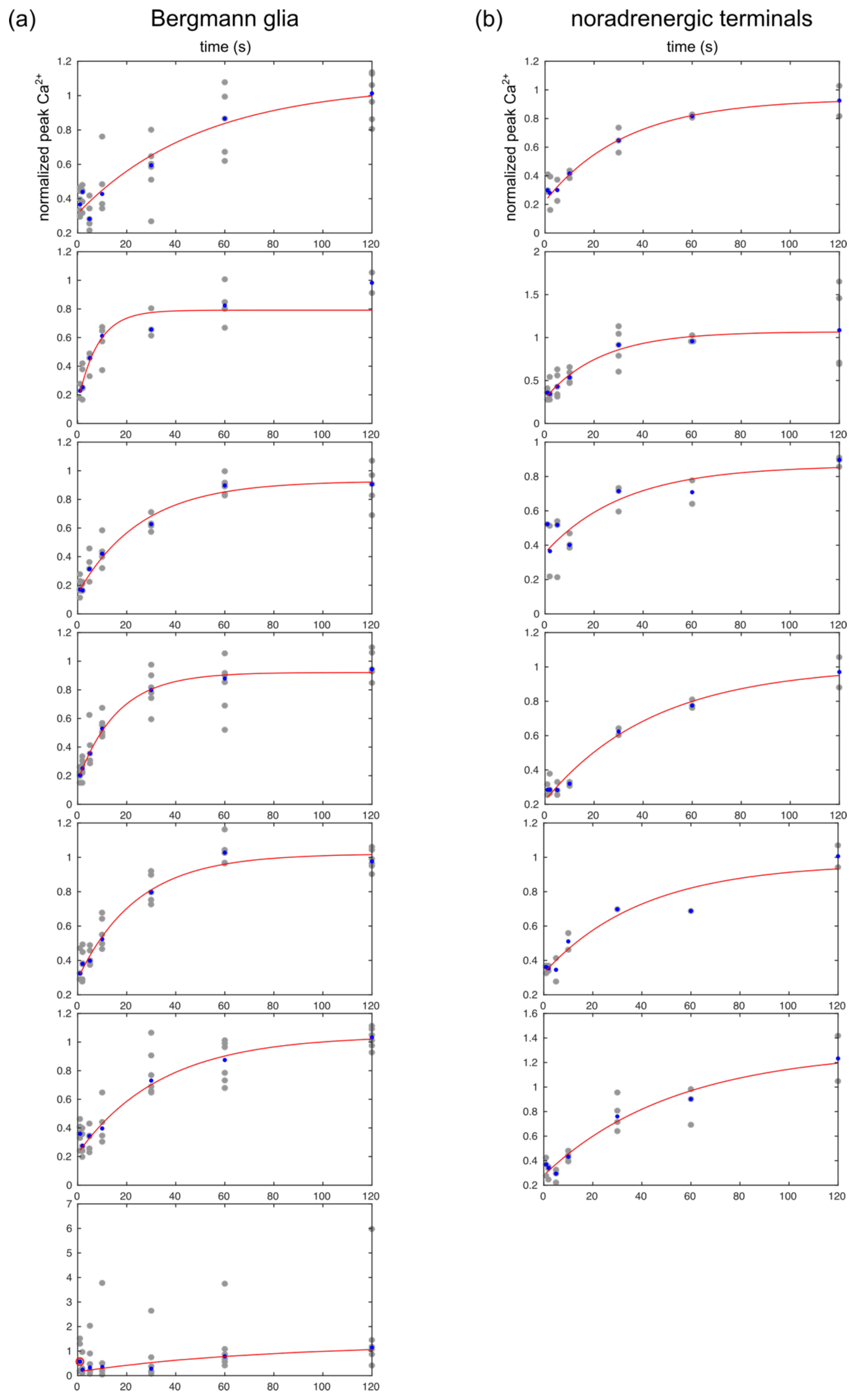


**Supplementary Figure 3**

**Exponential fitted plots generated for individual mice showing the varying extent of recovery of vigilance-dependent Bergmann glia and noradrenergic terminal $Ca^{2+}$ elevations between bouts of locomotion with varying lengths of intermediate pause.**
(a) Each plot represents the raw data (gray dots indicating individual repetitions), median values (blue dots) and fitted exponential curve (red trace) in a single mouse. *Slc1a3-CreER$^{T}$*;Ai95, *n* = 7 mice. In the last plot, the first normalized peak $Ca^{2+}$ value was an outlier and prevented the generation of a fitted curve using the same parameters as was used for all other experiments so it was excluded from generation of the fitted curve. (b) Same as in a for *Dbh-Cre*;*Lck-GCaMP6f* $^{flox}$, *n* = 6 mice.

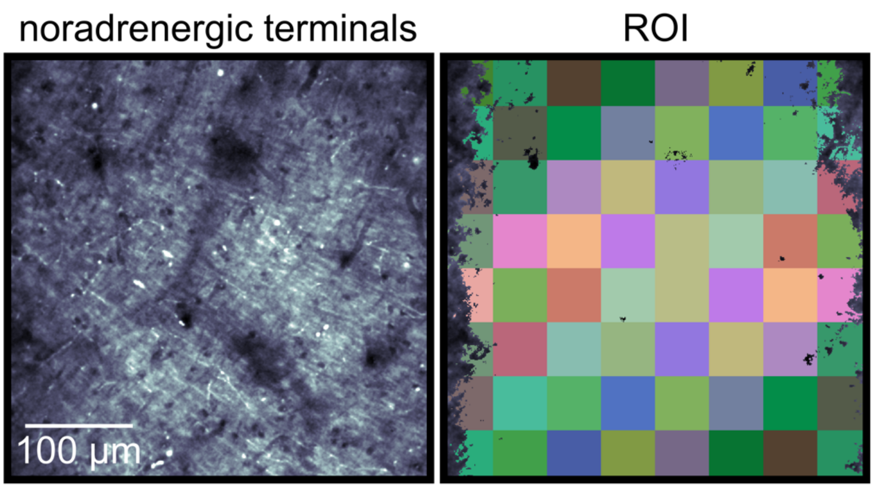


**Supplementary Figure 4**

**Representative field-of-view for a *Dbh-Cre*;Lck-GCaMP6f$^{flox}$ mouse.**

Image of *in vivo* GCaMP6f in noradrenergic terminal processes, tangential optical section ~60 µm beneath pial surface (left) and assigned ROIs (right).
